# Downgrading disease transmission risk estimates using terminal importations

**DOI:** 10.1101/265942

**Authors:** Spencer J Fox, Steven E Bellan, T Alex Perkins, Michael A Johansson, Lauren Ancel Meyers

## Abstract

As emerging and re-emerging infectious diseases like dengue, Ebola, chikungunya, and Zika threaten new populations worldwide, officials scramble to assess local severity and transmissibility, with little to no epidemiological history to draw upon. Standard methods for assessing autochthonous (local) transmission risk make either indirect estimates based on ecological suitability or direct estimates only after local cases accumulate. However, an overlooked source of epidemiological data that can meaningfully inform risk assessments prior to outbreak emergence is the absence of transmission by imported cases. Here, we present a method for updating *a priori* ecological estimates of transmission risk using real-time importation data. We demonstrate our method using Zika importation and transmission data from Texas in 2016, a high-risk region in the southern United States. Our updated risk estimates are lower than previously reported, with only six counties in Texas likely to sustain a Zika epidemic, and consistent with the number of autochthonous cases detected in 2017. Importation events can thereby provide critical, early insight into local transmission risks as infectious diseases expand their global reach.

## Introduction

The explosive emergence of Ebola in West Africa in 2014 and Zika in the Americas in 2016 caught the global health community by surprise. Officials scrambled not only to control the diseases at their source but also to anticipate and rapidly contain global transmission via infected travelers (1,2). The rate at which a newly introduced infectious disease spreads can vary enormously, depending on the physical and social environment. For example, serological surveys of dengue virus (DENV) exposure on either side of the Texas-Mexico border indicated far higher DENV exposure in the Mexican community despite virtually identical climatic conditions and even higher mosquito abundance in the Texan community (3).

Epidemiological risk assessment--estimating the severity and transmissibility of a threatening disease--can be vital to successful mitigation with limited resources. Historical outbreak data can provide invaluable insight into future epidemic risk. However, for a disease yet to arrive or that has just begun to spread, we necessarily borrow epidemiological data from other populations or related diseases, or to indirectly assess risk based on environmental suitability. For example, as the first importations of Zika virus (ZIKV) arrived in the US in 2016, early attempts to determine the likelihood and rate of local transmission relied primarily on dengue epidemiological data from regions with markedly different climatic and socioeconomic conditions (4–6).

These risk assessments provide information regarding the reproduction number of a disease (*R*_0_)---the expected number of secondary human infections resulting from a single human infection--- which provides a meaningful and predictive measure of local epidemiological risk. In a naive population, *R*_0_ indicates whether importations can potentially ignite local epidemics; if so, it also provides insight into the probability, magnitude, and speed of spread (7,8). Once a disease begins to spread, *R*_0_ can be directly estimated from early case data (9).

Here, we introduce a method for estimating *R*_0_ prior to an outbreak in populations that face the ongoing threat of infected travelers from affected regions. This approach was motivated by recent introductions of ZIKV into the continental US. As hundreds of cases arrived from affected regions throughout the Americas, officials sought to estimate risks of autochthonous (local) transmission and identify high risk regions in the southern US. However, given the novelty of ZIKV and the large proportion of ZIKV cases that go undetected, early estimates had high uncertainty (6,10,11). Our method harnesses importation data---individual cases that arrive in a naive location with or without subsequently infecting others---to update *a priori* estimates of *R*_0_, while explicitly modeling case reporting uncertainty. As a case study, we use the almost complete absence of secondary transmission following 298 importations of ZIKV into the state of Texas in 2016 and 2017 to reduce and narrow local estimates of *R*_0_.

## Methods

We used a two-step procedure to estimate the monthly *R*_0_ for each of the 254 Texas counties (hereafter county-month *R*_0_): (1) estimate *a priori* county-month *R*_0_ distributions using published ecological models of ZIKV transmission (4,6), and (2) using these as Bayesian priors, generate posterior *R*_0_ distributions based on reported importations and subsequent local transmission.

### Data

We analyzed all ZIKV importations into Texas from January 2016 to September of 2017, including the county and notification date. County-level purchasing power parity (PPP) in US dollars (12); daily temperature data at a 5 km x 5 km resolution for 2016-2017 and historical averages from 1960-1990 (13,14) were also used as inputs to the transmission risk model. For each county and month, we averaged daily temperatures across all 5 km x 5 km grid cells whose center fell within the county; we aggregated 5 km x 5 km mosquito (*Aedes aegypti*) occurrence probabilities similarly (15). Data available doi:10.18738/T8/HYZ53B.

In all, six mosquito-borne, autochthonous cases of ZIKV were reported in Texas in 2016 and two were reported in 2017 (25). For updating *R*_0_ estimates, we analyzed 2016 data and assumed that two autochthonous cases were detected in Cameron County--one in November and one in December 2016; we excluded four nearby cases discovered during the November follow-up investigation, because our model does not incorporate active surveillance. As sensitivity analyses, we re-estimated *R*_0_ assuming that no cases were detected and that all six cases were detected (Fig S7). For validating our estimates, we analyzed 2017 data and considered only one of the two reported autochthonous cases, as the second case occurred outside the timeline of our 2017 importation data.

### A priori county-month R_0_ estimates

Following Perkins et al (6), we estimated *R*_0_ using the Ross-Macdonald temperature-dependent formulation:

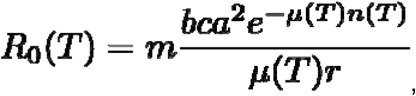

with parameters as defined in Table 1. We calculated relative abundance of the ZIKV vector based on *Ae. aegypti* occurrence probabilities as-ln(1-occurrence probability), and interpret this as a relative (rather than absolute) abundance, which is sufficient for our *R*_0_ estimation (6). We derived *a priori* county-month *R*_0_ distributions by drawing 1,000 Monte Carlo samples from each underlying parameter distribution, with the appropriate county and month data. Finally, we fit gamma distributions to each probability distribution for use as an informative priors.

**Table 1:** Parameters of prior *R*_0_ estimates.

| Parameter | Description | Distribution | Value (CI) | Citation |
| --- | --- | --- | --- | --- |
| $b$ | Mosquito-to-human transmission probability | Constant | 0.4 | (16) |
| $c/r$ | Human-to-mosquito transmission probability times the duration of human infectiousness | Constant | 3.5 | (17) |
| $a$ | Mosquito biting rate | Constant | 0.67 | (18) |
| $\mu(T)$ | Mosquito daily mortality rate | Non-parametric GAM | 0.115<sup>1</sup> | (19,20) |
| $n(T)$ | Extrinsic incubation period in Mosquitoes | Exponential | 6.1 (3.4, 9.9)<sup>2</sup> | (21) |
| $m$ | Economic mosquito-human contact factor | Monotonic decreasing SCAM | Fit to monte carlo samples | (6) |

1. Fit to data from mark-recapture study occurring between 20-34 C 2. At 30 C

### Autochthonous transmission likelihood

Following (22), we developed a likelihood function describing the expected outbreak size following an importation. We assumed that the secondary case distribution for each infected is negative binomial with mean *R*_0_, and dispersion parameter, *k*. Assuming all cases are detected, the probability of an outbreak of chain size, *j*, is given by:

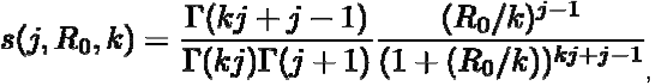

where (n) = (n-1)!. However, not all cases are detected and the imported index case is always detected and correctly classified as an importation, so the probability of detecting a chain of size, *j*, from a given importation is given by:

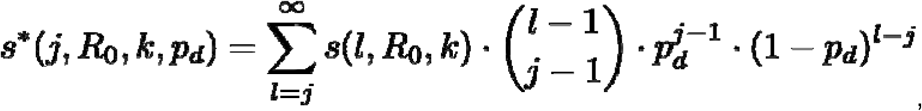

where *p_d_* is the case detection probability. Importantly, this allows for local, undetected cases.

We take the product of all likelihoods for each imported case as

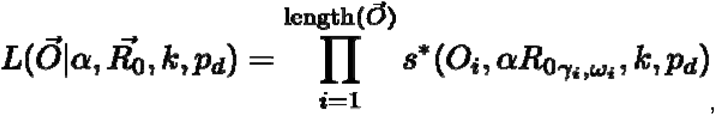

where the vector, *O*, contains the observed outbreak sizes for each importation (terminal importations have an outbreak size of one), denotes the county ()-month () *R*_0_ for the location and time that the importation occurred, and α is a statewide scaling factor applied to each. The introduction of the state-wide scaling factor allows for localized importations to inform statewide estimates, but assumes that biases in the *a priori R*_0_ estimation procedure are constant across counties and months. Details of simulations and validation of the likelihood can be found in supplemental section I (Fig S1).

### Estimating the dispersion parameter

The negative binomial dispersion parameter governs the variability in secondary cases following each importation, with values near zero meaning that most importations yield few or no cases while a few “superspreaders” produce many. We assume that ZIKV secondary case distributions resemble that of dengue virus (DENV) (23). Padmanabha *et al.* describe the relationship between regional *R*_0_ and the percentage of DENV cases generating over 20 secondary infections (*p_20_*), as *R*_0_ = 0.63 x 100 x (*p_20_*) + 0.58. We assumed that *p_20_* = 1e-8 for *R*_0_<0.58, and found that a single dispersion parameter captures this relationship for all *R*_0_ values and thus used k=0.12 for all analyses (Fig S2).

### Updating posterior R_0_ estimates

We estimated posterior distributions for α, and each county-month *R*_0_ for each day with a new importation between January 2016 and January 2017. We assumed a uniform prior for α of U∼(0,2), and used a blocked Gibbs sampling algorithm of MCMC. For each MCMC step we provide the detected imported cases to date and propose each county-month *R*_0_, a single α, and a *p_d_*. County-month *R*_0_ proposals were normally distributed around the previous sample with standard deviation of 0.1, α proposals were distributed U∼(0,2), and we used a previously estimated distribution for the reporting rate, *p_d_* ∼ N(5.74%, sd=1.49%), which we assumed to not vary spatiotemporally (24). We used the Metropolis-Hastings probability to accept or reject proposals. Our chains consisted of 200,000 samples with a burn-in duration of 100,000; thinning every 10 steps. Further algorithmic details and code are available on Github (https://github.com/sjfox/rnot_updater).

### Validating posterior county-month R_0_ estimates

We derived the expected number of autochthonous cases from the importations data through September of 2017 (at that time, the most recent importation was detected in mid-May) and compared the estimates to the actual reported autochthonous cases. We integrated uncertainty into our estimates by sampling from the posterior county-month *R*_0_ distributions and simulating outbreaks accordingly (full details in supplemental section II).

**Figure 1:**
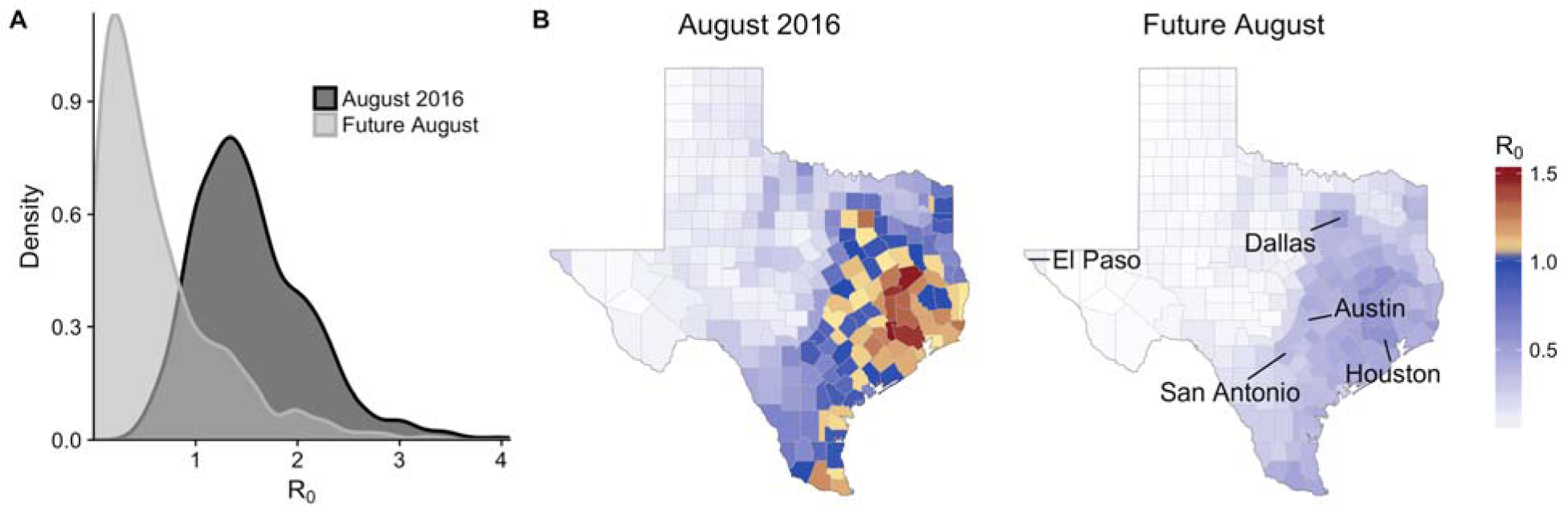
*R*_0_ updating using importation data. Consider a hypothetical scenario in which the first 15 terminal ZIKV importations into Texas arrive in Harris County (which includes Houston) during August 2016. **(A)** Estimated Harris County *R*_0_ for August 2016 *a priori* (dark grey) and after accounting for the 15 (light grey) terminal importations (Future August). **(B)** Median *R*_0_ estimates for August before (August 2016) and following (Future August) the importation-based update.

## Results

### Importation-based updates of transmission risk

Hypothetically, suppose that the first 15 imported cases of Zika into Texas arrived in August into Harris County (which contains Houston) without any detected autochthonous transmission. Prior to these importations, environmental suitability models yielded a relatively high local risk estimate with median Harris county *R*_0_ above the epidemic threshold of one (Figure 1A - dark grey). The lack of secondary cases following all 15 importations suggests that *R*_0_ may be lower. Indeed, our updated estimates suggest that the Harris county *R*_0_ is likely below one (Figure 1A - light grey). Our method leverages such county-level importation data to update *R*_0_ estimates throughout the state (via a scaling factor), based on the assumption that any *a priori* biases will be similar across counties (Figure 1B).

### Baseline importation and transmission risks in Texas

Prior to making importation-based updates, our initial median estimates of *R*_0_ across Texas’ 254 counties in 2016 range from approximately 0 to 1.5 throughout the year with July and August having the highest transmission risk (Figure 2A). Throughout the manuscript, we conduct a one-sided test at a 1% significance level and thus consider counties with 99 percentiles (upper bounds) that include one to be at risk for an epidemic (*R*_0_ > 1). Initial upper bound estimates reach as high as three, and 119 (47%) of Texas counties are expected to be at risk of a local outbreak in at least one month of the year (Figure 2A, S2). When we considered historic average temperatures rather than 2016 temperatures, the projected 2017 risks were consistently lower, with the largest differences occurring during the unseasonably warm 2017 winter (Fig S4). Case importations peaked in July, August, and September of 2016, with 164 (55%) of the 298 total 2016 importations arriving then (Figure 2B). The few detected autochthonous cases occurred in November and December, when expected risk was relatively low but not negligible.

**Figure 2:**
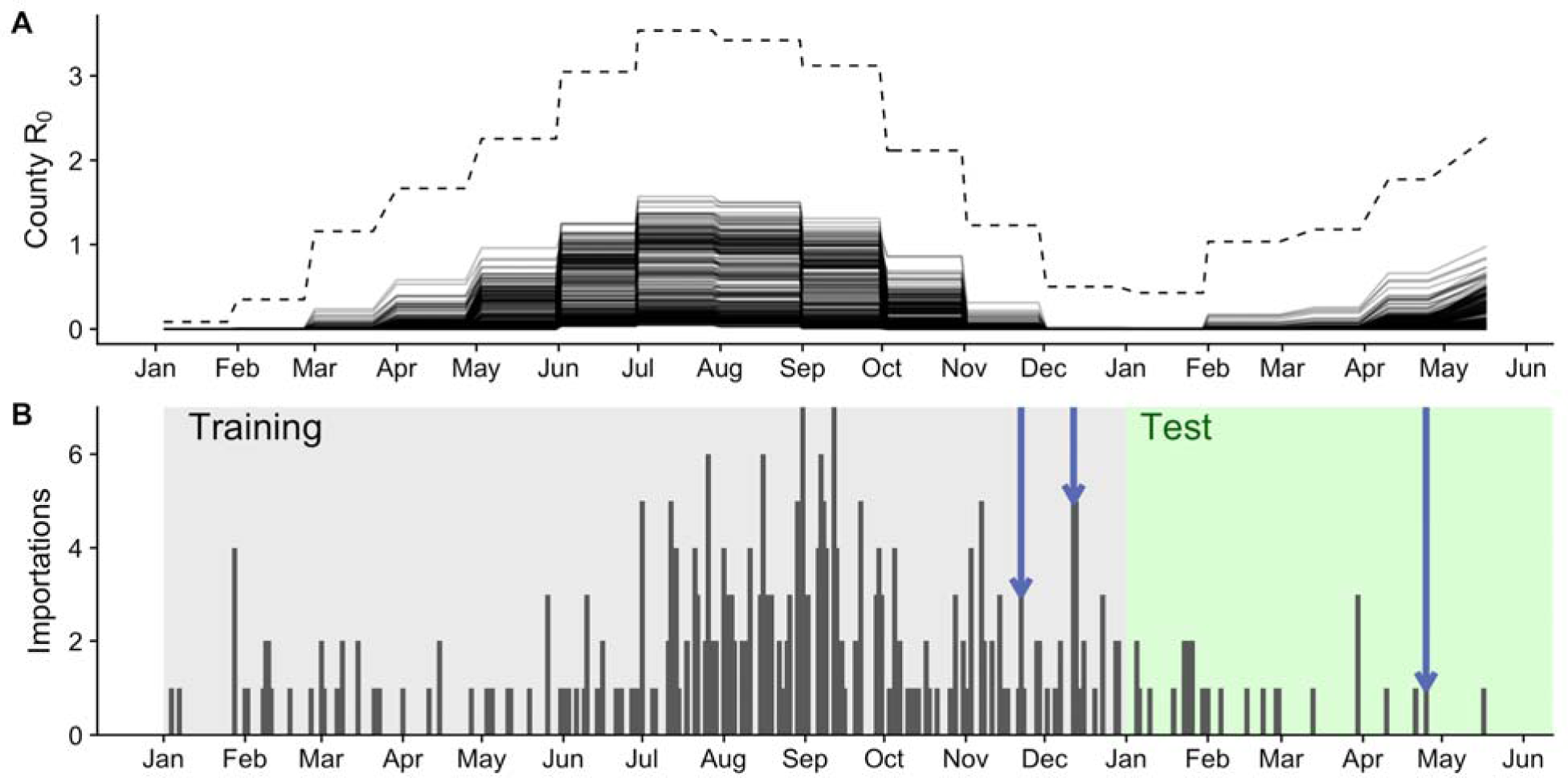
Texas importations and baseline transmission risk estimates for 2016-17. **(A)** Initial ZIKA *R*_0_ estimates using ecological risk models parameterized with actual 2016-2017 temperatures. Each solid line shows median values for one of Texas’ 254 counties. Dashed line shows the highest upper bound (99th percentile) across all counties. **(B)** Daily ZIKV importations into Texas. Blue arrows indicate importations that produced detected autochthonous transmission; shading indicates training (2016) and testing (2017) periods.

### Updated transmission risks in Texas

Based on all importations and autochthonous cases that occurred in Texas prior to January 2017, we estimate that all Texas counties have a median posterior *R*_0_ below one (Fig 3). Median estimates range from 0 to 0.29; upper-bound estimates range from 0 to 1.12, with only six (5%) of the original 119 high-risk counties maintaining epidemic potential (Fig S5). When we assume historic averages rather than 2016 temperatures, we obtain similar results (Fig S6).

In a sensitivity analysis that assumes ∼20 times more *undetected* importations, we found that the estimated risks decreased further (Fig S7). We also varied the number of detected autochthonous cases in November: as they decrease from one to zero, the estimated risks decrease considerably; as they increase to five, estimated risks increase, with 83 counties becoming at risk for a local outbreak (Fig S7).

**Figure 3:**
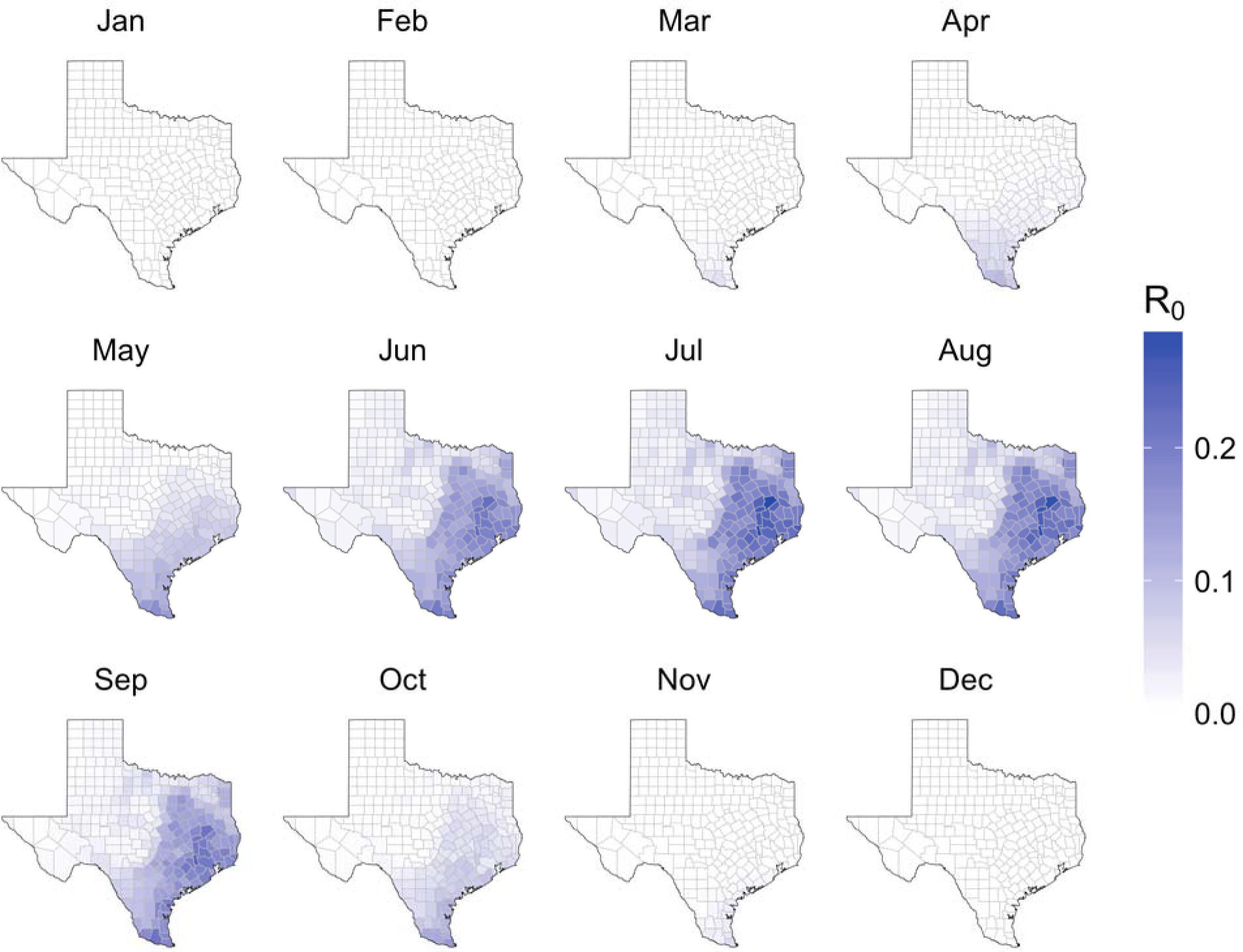
Posterior median county *R*_0_ estimates for Texas, based on ZIKV importations through January 2017. This assumes that all importations were terminal except for a two autochthonous cases detected in Cameron County in late 2016.

Importation events had variable impacts on the posterior estimates, depending on their timing and location (Fig 4). Terminal importations early in the year, when *a priori R*_0_ estimates were low, had little effect; those arriving in the summer months, when high *a priori R*_0_ estimates suggested that transmission should have occurred, led to sharp decreases and a shrinking confidence interval. By early November, the median α decreased from 1.0 to 0.06 with a narrow 95% CI of 0.002-0.30. However, the two secondary transmission events detected in November and December increase *R*_0_ estimates and widen the confidence intervals. Incorporating all data up to January 2017, our best estimate is that *R*_0_ values across the state are roughly one fifth the original estimates (median: 0.19, 95% CI: 0.05-0.48).

**Figure 4:**
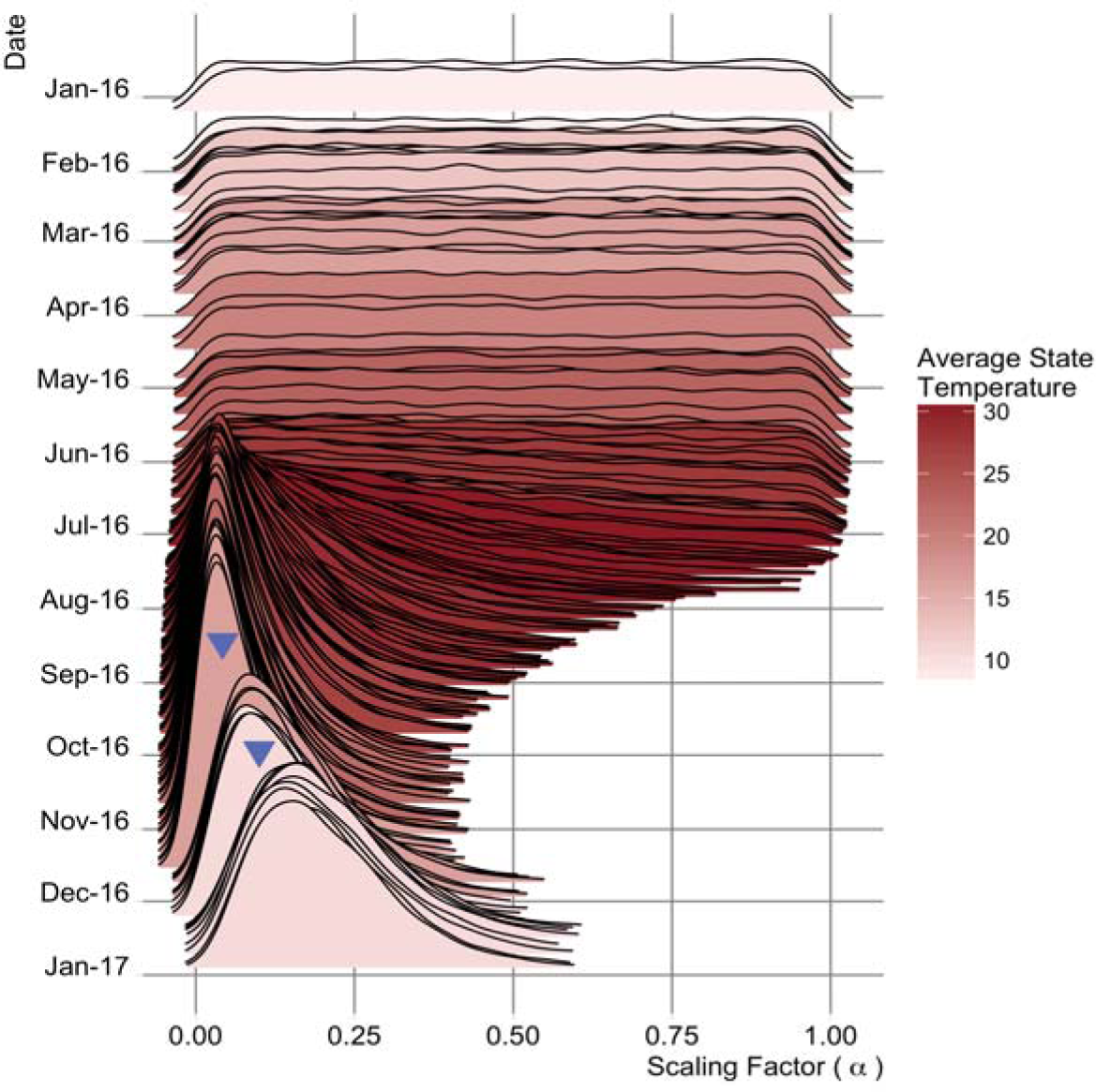
Evolving posterior distribution of statewide scaling factor for *R*_0_. Zika importations, both with and without subsequent detected autochthonous transmission, provide insight into local transmission potential, via a statewide scaling factor, **α**. This shows the posterior distributions of **α**, for each day of 2016 that had at least one imported case. Median estimates reach a minimum in early November, just before the detected autochthonous transmission events (upside-down blue triangles). Red shading indicates the average statewide monthly temperature. Note: the scaling factor is never less than zero.

### Expected autochthonous transmission in Texas

We use transmission risk estimates based on importations through December 2016 to estimate the number of autochthonous cases we would expect to detect in Texas in 2017. Assuming first that only the reported importations occurred in 2017 (26 total), we estimate that there should have been 0.08 (95%CI: 0-1) detected autochthonous cases in the state; assuming that many importations went undetected, according to the reporting probability (26 / *p_d_* ≈ 453 total), we estimate 1.3 (95% CI: 0-7) detected autochthonous cases. These estimates are consistent with the single autochthonous case detected in Texas in 2017 (Fig 5).

**Figure 5:**
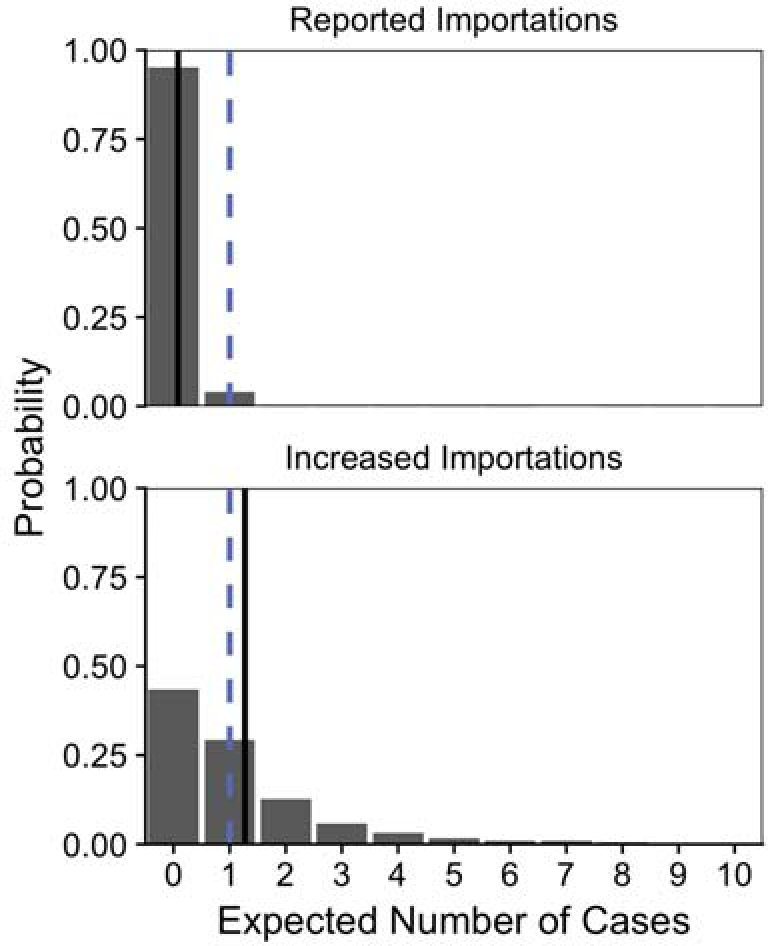
Expected autochthonous cases in 2017, assuming revised county *R*_0_ estimates and reported importations through September 2017. The probability distributions are built from 10,000 simulations, each randomly drawing from the *R*_0_ posterior distributions. The dashed blue line indicates the actual number of detected autochthonous cases in state (one), and the solid black lines indicate means for the baseline importation scenario, in which only the reported importations occurred (top) and the increased importation scenario, in which a large fraction of importations went undetected (bottom).

## Discussion

The global expansion of ZIKV was declared a Public Health Emergency of International Concern in February 2016, and caused more than 565,000 confirmed or probable cases and over 3,352 documented cases of congenital Zika syndrome. Although it is receding in most regions of the world, ecological risk assessments suggest that previously unaffected or minimally affected areas may remain at risk for future emergence, including parts of Asia and South America (26–28). Differentiating regions that can sustain a ZIKV epidemic (*R*_0_>1) from those that cannot is vital to effective planning and resource allocation for future preparedness plans. To address this challenge, we have developed a simple method for refining uncertain risk assessments with readily available data on disease importations.

We applied the method to update ZIKV *R*_0_ estimates for each of the 254 counties in Texas, and found that only six counties have non-negligible probabilities of sustained local transmission. This is a substantial downgrade in expected risk, given that 43% of the 254 counties were previously thought to be vulnerable to ZIKV outbreaks. These estimates suggest that there should have been roughly one detected case of locally acquired ZIKV between January and September of 2017, closely corresponding to the single transmission event actually detected in Cameron County in July 2017 (Figure 5). Our sensitivity analysis suggests that, if we underestimated case-reporting in November, 77 additional counties have non-negligible but low risks of summer outbreaks. Given comparable importation and climatic data, this approach could readily update ZIKV transmission risk estimates for all counties in the continental US and elsewhere.

Our estimation method relies on several simplifying assumptions. We assumed that the shape of the secondary case distribution resembles that of dengue. Although we have no evidence to the contrary, this should be updated as ZIKV-specific estimates become available (23). We also assumed that transmission is equally likely from imported and locally acquired cases. Imported cases may be less infectious than locally acquired cases for two reasons, leading us to underestimate local transmission risks. First, they may be more likely to receive care or education that limits subsequent transmission, although most ZIKV cases are inapparent or mild, and do not require medical care (11); second, if they arrive already infectious, their local infectious periods may be shorter than those of autochthonous cases. Next, we treat all importations as independent. However, spatiotemporal heterogeneity in case detection probabilities or clustering of cases (e.g., testing of travel companions) could bias risk estimates. Furthermore, when secondary clusters are detected, we assume they share a transmission tree stemming from a single detected importation. In fact, the low ZIKV detection rate suggests that both primary importations and secondary cases are likely to be missed. If the detection rates are roughly similar, our results hold. When we assume, in sensitivity analysis, that importations are detected at higher rates than secondary cases, then the resulting risk estimates will be higher; when we assume the reverse, they are lower. The additional assumption, that clusters are epidemiologically connected, seems reasonable for the small contained outbreaks detected in Texas, but may not be appropriate for importation-fueled arbovirus outbreaks in Florida, for example. In such cases, molecular data might enable estimation of transmission clusters (32,33). We also rely on informative Bayesian priors and a statewide scaling factor, which allows us to use local importations to inform risk estimates elsewhere, but implies that our prior county-month transmission risk estimates are correct relative to each other. Given additional importation data, we could potentially estimate each county-month *R*_0_ independently. Finally, we do not consider possibility of sexual transmission of ZIKV. While sexual transmission has occurred and may be important for specific populations (29), we assumed that mosquito-borne transmission is the dominant mode of infection.

During the height of the ZIKV threat, public health agencies in the US rapidly implemented both preventative measures (e.g., vector control and educational campaigns) and response measures (e.g. laboratory testing and epidemic trigger plans), particularly in high risk southern states. Decision makers sought to identify and narrow the spatiotemporal scope of outbreak risk to enable targeted responses, efficiently allocate resources, and avoid false alarms (10,30). Our method facilitates such rapid, real-time geographic risk estimation from typical early outbreak data, and suggests that only 3% of the Texas population is at risk for a local outbreak. Critically, we can conclude neither that all initial ecological risk assessments for ZIKV will overestimate risk, although this seems to be the case for ZIKV in Texas, nor that public health preparations and interventions for ZIKV are no longer necessary in Texas or the southern US. Rather, our results suggest that sustained ZIKV outbreaks are unlikely, but not impossible, and provide more robust and localized estimates of ZIKV risk that can inform more targeted surveillance and reactions to future ZIKV importations.

This framework can be applied to update any *R*_0_ estimates using importation data, regardless of the *a priori* method of estimation. For example, a new approach combining epidemiological and molecular analyses suggests that transmission risk in Florida is subcritical (i.e., *R*_0_ < 1) (31,32). Given that Florida experienced thousands of introductions, only a few of which sparked large outbreaks, coupling such outbreak-driven estimation with our terminal importation method may provide a powerful real-time risk assessment framework for exploiting all available data.

We presented a simple and rational method for continuously updating transmission risk estimates for populations experiencing infectious disease importations, with or without secondary transmission. As we demonstrated for ZIKV in Texas, large numbers of terminal importations can profoundly lower both estimated risks of transmission and uncertainty in prior estimates, particularly those derived from ecological suitability or other models that borrow inputs from related pathogens in other parts of the world. Although the threat of ZIKV emergence in the continental US motivated this study, this new framework can be widely applied to improve transmission risk assessments when a disease newly threatens a population via regular introductions with minimal secondary transmission. For example, importation-fueled MERS-CoV transmission risk, or highly pathogenic avian influenza (34,35). This method can also be used to assess disease transmission risk during elimination scenarios, such as assessing risk for measles transmission in vaccinated populations or malaria in non-endemic regions (36,37).

## Acknowledgements

The authors would like to acknowledge Texas DSHS for providing the county-level ZIKV importation data used in this analysis. SJF and LAM would like to acknowledge funding from the Models of Infectious Disease Agent Study (MIDAS) program grant number U01GM087719; SEB received support from a National Institute of Health National Institute of Allergy and Infectious Diseases grant (K01AI125830); MAJ received partial support from the MIDAS program (Cooperative Agreement number 1U54GM088558); and TAP received support from the National Science Foundation (DEB 1641130) and the Defense Advanced Research Projects Agency (D16AP00114). Finally, we would like to acknowledge the Texas Advanced Computing Center (TACC) at The University of Texas at Austin for providing high performance computing resources that have contributed to the research results reported within this paper. URL: http://www.tacc.utexas.edu.

## Author Contributions

SEB, LAM, and SJF developed the conceptual modelling framework. SJF completed the analyses and created the figures. SJF, TAP, and MAJ reviewed published literature. All authors contributed to the interpretation and presentation of results. SJF wrote the first draft, and all authors contributed to writing and approval of the final report.

## Data and code availability

Texas county-level importation data has not been given approval for public release, but all other necessary data can be found online (http://dx.doi.org/10.18738/T8/HYZ53B) and the code can be found on github (https://github.com/sjfox/rnot_updater). These data and code repositories contain fake importation data that can be used to fully understand the statistical methods presented here. Please contact Texas DSHS directly for access to the real county-level importation data used in the analysis.

